# Identification of minor alleles associated with reduced lodging in tef (*Eragrostis tef*)

**DOI:** 10.1101/2022.03.17.484745

**Authors:** Shiran Ben-Zeev, Timo Hellwig, Muluken Demeile, Vered Barak, Sasha Vorobyova, Sariel Hübner, Yehoshua Saranga

## Abstract

**Rational:** Underutilized species that are not widely cultivated (known as orphan crops) present opportunities to increase crop diversity and food security. Tef [*Eragrostis tef* (Zucc.) Trotter] is known for its high-quality grain and forage. Root-borne lodging is a major devastating problem in tef cultivation, leading to large economic losses and limiting its widespread adoption.

**Objective:** The aim of this study was to identify genomic regions that are associated with tef lodging.

**Methods:** A tef diversity panel (TDP-300) comprised of 297 lines was assembled, genotyped, and phenotyped across 4 field environments. This unique panel, the first of its kind in tef, has the potential to facilitate tef research and breeding.

**Results:** Genome-wide association study identified 29 sites associated with lodging; in all cases with a minor allele conferring reduced lodging. The eleven sites of prime interest were located in or near genes, 5 of them with a putative role, of which 3 were found to be involved root development.

**Conclusions:** The identification of lodging-related sites in the current study may advance understanding of the mechanisms underlying tef lodging and crop improvement. The identification of genes related to root development support the importance of root traits in tef lodging, which should be targeted in future breeding.

## Introduction

Sustaining agricultural productivity and food security is becoming ever more challenging for many reasons, including climate change (Rosenzweig *et al*. 2014), lack of diversity (Renard and Tilman 2019; Tadele 2019), diminishing soil and water resources (Wuepper *et al*. 2020), increasing human population (Tilman *et al*. 2011) and changing global consumer demands (Khoury *et al*. 2014). These dramatic changes are affecting the world’s major crops - maize, wheat and rice (Zhang *et al*. 2021). A viable strategy to enhance agricultural productivity and food security is to increase crop diversity (Huang *et al*. 2002), with a particular emphasis on resilience to challenging environmental conditions such as waterlogging, drought and salinity (Smith *et al*. 1989; Renard and Tilman 2019; Girija *et al*. 2021). The diversification of crops grown for human and animal consumption is important ecologically as well as nutritionally (Khoury *et al*. 2014) and it is even more salient in light of the current climate change (Kamenya *et al*. 2021).

Underutilized species that are not widely cultivated (known as orphan crops) present opportunities to increase crop diversity. The reasons for their minor importance in global agriculture have been debated (Tadele 2019; Allaby 2021; Milla and Osborne 2021), but there is consensus on their potential to improve food production and security. Some information on growing conditions, genetics and ecology is available for a number of orphan crops (Mabhaudhi *et al*. 2019). Combining the existing information with genomics and genetics could help convert them into large-scale crops, thus benefiting from both their diversity and their resilience (Varshney *et al*. 2012; Dawson *et al*. 2019). An important tool for the introduction of a new crop into modern agriculture, and to support its breeding and development, is a genomic database (Brook *et al*. 2008). Genomic insights into the control of various traits have been proposed as a target for orphan crops (Varshney et al. 2012). Genome-wide association studies (GWAS) have long been used to expose the genetic basis of traits (Ikegawa 2012), and the use of genomic data derived from genotyping-by-sequencing (GBS) assays for GWAS is an accepted approach (Bellucci *et al*. 2015). Recently, GWAS have been applied to several African orphan crops: fonio millet (*Digitaria exilis*) (Abrouk *et al*. 2020), finger millet (*Eleusine coracana*) (Puranik *et al*. 2020) and pigeon pea (*Cajanus cajan*) (Zhao *et al*. 2020). The key role of orphan crops in areas that are unsuitable for common major crops suggests their tremendous potential for introduction into challenging environments. In addition, orphan crops contain genes that may help develop similar resilience in major crops (Jamnadass *et al*. 2020; Kamenya *et al*. 2021).

Tef [*Eragrostis tef* (Zucc.) Trotter] is an allotetraploid C4 (2n = 20) cereal crop with a genome size of 622 Mbp (VanBuren *et al*. 2020), mostly grown in Ethiopia where it was presumably domesticated (D’Andrea 2008). Tef grains are gluten-free, and contain high amounts of nutritious compounds and all nine essential amino acids (Reda 2014; Shumoy *et al*. 2018; Kebede 2021); it is therefore commonly referred to as a “superfood”. In addition, tef is an excellent forage grass (Ditsch 2007; Matthew Davidson 2018). Tef grows in a wide range of environments, from sea level to over 2000 m elevation, with seasonal precipitation of 200 to over 2000 mm, and from drought-prone areas and saline soils to water logging conditions (Ketema 1983a, 1993; Assefa *et al*. 2013). These characteristics facilitate the interest in adopting tef in countries outside the Horn of Africa (Miller 2008; Norberg *et al*. 2009; Brown 2017; Ben-Zeev *et al*. 2018).

Adapting a traditional crop, such as tef, to modern, highly mechanized agriculture requires reevaluation and calibration of agronomic practices in tandem with genomic studies and breeding. The introduction of tef as a new crop to irrigated Mediterranean conditions presents many challenges, at each step of its growth, some of which have been targeted in our recent studies (Ben-Zeev, Kerzner, *et al*. 2020; Ben-Zeev, Rabinovitz, *et al*. 2020). The relatively minor attention paid to tef in research, as well as the lack of a reference genome and diversity panel, has hindered studies on the genomic basis of key traits in this crop plant (Girija *et al*. 2021). Nevertheless, some progress has been made in these aspects over the last decades. AFLP (Bai *et al*. 1999), RFLP (Zhang *et al*. 2001) SSR (Yu et al. 2006, 2007; Zeid et al. 2011) markers were used to map QTLs for various traits onto linkage groups. Two tef reference genomes (Cannarozzi *et al*. 2014; VanBuren *et al*. 2020) were recently published, the latter identifying exceptional subgenome divergence. A most recent study used the published reference genome to identify genomic regions associated with tef grain nutritional traits (Ereful *et al*. 2022). These studies are opening new avenues for further exploration of tef’s genomic potential. Recently published reviews have called for genomic studies specifically in tef, showcasing their contribution to other cereals (Girija *et al*. 2021; Numan *et al*. 2021).

Plant lodging, defined as the permanent displacement of the culm from its vertical position, is the most devastating problem in tef cultivation. Lodging can be divided into stem lodging, in which the stem bends or breaks, and root lodging, in which the angle between the stem and the soil is a changed due to root disanchoring (Berry *et al*. 2004), the latter being the prime cause of tef lodging(Ben-Zeev, Rabinovitz, *et al*. 2020). Various studies estimated that tef lodging cause 30– 50% yield losses (Ketema 1997; Van Delden *et al*. 2010; Zeid *et al*. 2010), reduce grain and feed quality (Ketema 1983b; Van Delden *et al*. 2012; Reda 2014) and lead to large economic penalties. The damage caused by lodging is exacerbated in modern mechanized farming due to the difficulty in harvesting lodging plants. Irrigation, required for tef cultivation under semiarid and arid conditions (Ben-Zeev *et al*. 2018) and mostly applied via sprinklers or pivots, further intensifies lodging due to increased canopy weight while the soil is lubricated and the plant’s anchorage loosened. Lodging often occurs on a plot or field level - meaning that a single plant or group of plants which start lodging have a literal “domino” effect (Fig. 1).

**Figure 1.**
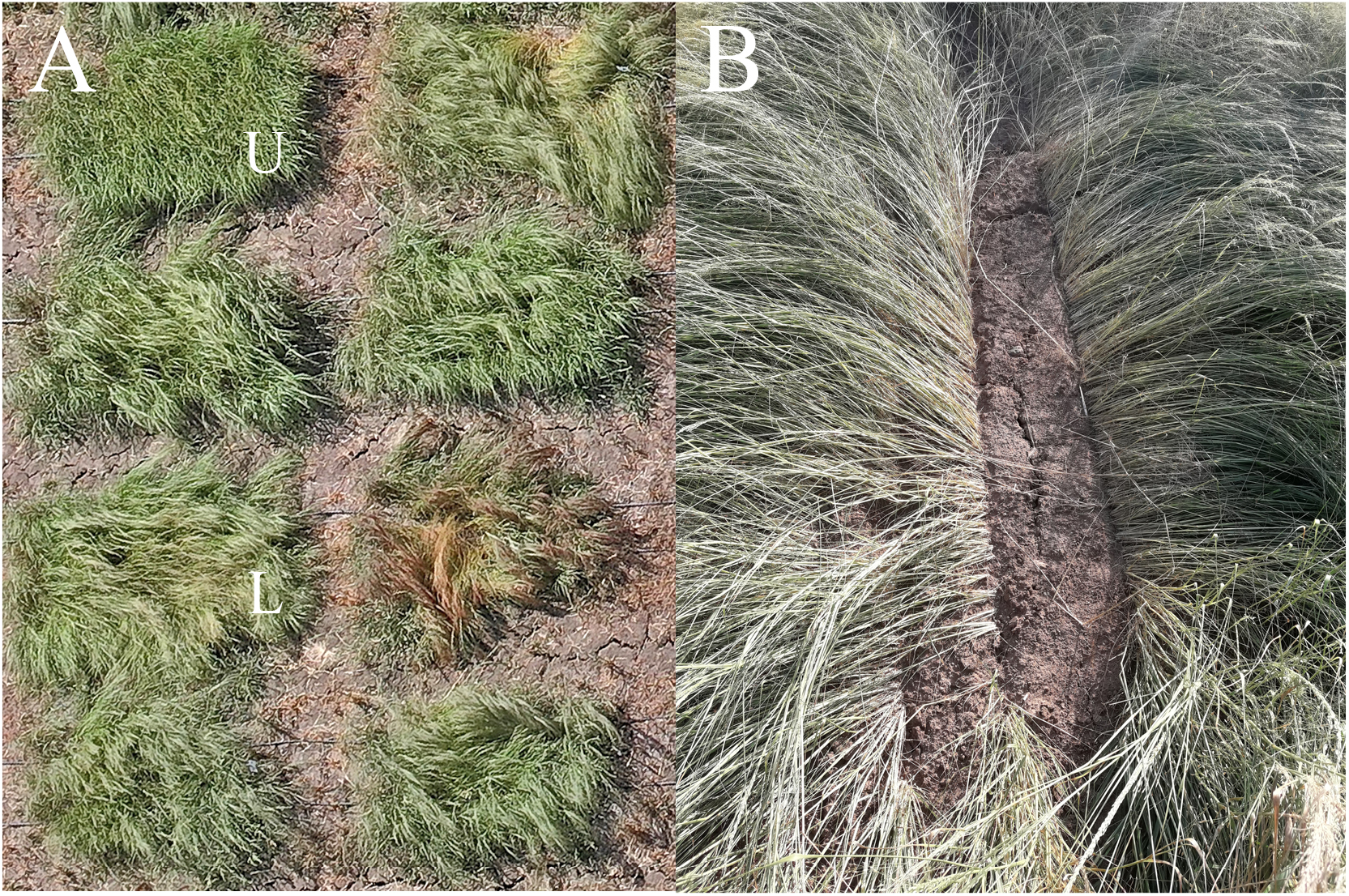
(a) drone photograph of lodging (L) and upright (U) experimental plot at 9 weeks after emergence the 2021 experiment, and (b) close up photo of a lodging plants.

Lodging is a dynamic and detrimental phenomenon (Berry *et al*. 2000, 2004; Van Delden *et al*. 2010; Blösch *et al*. 2020), which is most likely to be affected by various characteristics and genes at different developmental stages. Like most other traits, it is affected by environment (E), management (M), genotype (G) and their interaction (G x E x M). The effects of select management practices on lodging have been reported (Paff and Asseng 2018 and references therein). Semi-dwarfism, as a basis for reduced lodging, has been targeted via mutagenesis (Zhu et al. 2012; Jost et al. 2015), as well as by multiparental (Abraha et al. 2017) and biparental populations (Bekana and Assefa 2021). In the current paper, we report on the identification of genomic regions associated with plant lodging, possibly the most devastating and challenging tef phenotype, thus paving the way towards breeding for lodging resistance.

## Materials and Methods

### Diversity panel assembly and genotyping

A set of ca. 400 tef accessions, collected by the USDA in Ethiopia during the 1950s and 60s, and stored since the early 1990s in the Israeli Gene Bank (Agricultural Research Organization, Volcani Center) was propagated and stabilized via single-plant selection (Ben-Zeev *et al*. 2018). DNA was isolated from leaf tissue at the seedling stage using a Nucleospin II kit (Mecherey Nagel, Germany). Sequencing was conducted at the Australian Genome Research Facility (AGRF), using a ddRAD-based library-preparation protocol. DNA was digested using restriction enzymes Pstl and Nlall, ligated to adaptors, amplified, and sequenced. Reads were tested for quality, aligned to the tef reference genome (VanBuren *et al*. 2020), and variants were called using the Stacks V2.3d pipeline (Catchen *et al*. 2011).

In the subsequent filtering steps, only biallelic SNPs were retained. Genotype quality was set to a minimum of 15. A minimum of 5 reads was set to call a genotype, where heterozygous genotypes with read depth (DP) < 30 were converted to missing data. Minimum and maximum mean depth over individuals were set to 5 and 20, respectively, based on the depth distribution (Fig. S1a). The lines were highly homozygous, owing to tef’s self-pollinating nature and the single-plant-selection origin of the samples. Due to the high homozygosity and the abundance of rare alleles, minor allele frequency (MAF) was set to >1%. Sites with >30% missing data were removed. Individual lines with >1% heterozygous loci and/or >30% missing genotypes were excluded from the data set. Individuals were filtered based on identity by state (IBS) using four internal controls (lines doubled in sequencing). The final variant calling file (VCF) included 28,837 polymorphic sites and 297 lines, hereafter termed tef diversity panel 300 (TDP-300).

### Population structure

Linkage disequilibrium (LD) pruning was conducted with Plink (Purcell *et al*. 2007). Windows of 20 kbp were shifted by 5 variants at each step and variants in one window were removed if they exhibited r^2^ > 0.5. Population stratification was explored using sparse non-negative matrix factorization (sNMF) with the LD-pruned SNP set algorithm as implemented in the R package ‘LEA’ (Frichot and François 2015). Because the regularization parameter (α) can have a large influence on the outcome of the analysis’ incomparably small data sets (Frichot *et al*. 2014), the optimum α was determined by testing a range of α values. The α value with the minimum mean cross-entropy error was used for the final analysis. The final analysis was performed with the following parameters: α = 260, K ranging from 1 through 30, 10 independent repetitions, 5,000 iterations, and a tolerance error of 10^-6^. Cross-entropy was calculated with 10% masked genotypes. Model-free approaches such as sNMF do not depend on the explicit assumption of Hardy–Weinberg equilibrium and therefore, they are appropriate for both inbreeding and outbreeding species (Frichot *et al*. 2014).

A neighbor-joining tree was created based on Prevosti’s distance (Prevosti *et al*. 1975) and 100 bootstrap values to estimate the node support. Principal coordinate analysis (PCoA) was performed using GENPOFAD genetic distances (Joly *et al*. 2015; R Core Team 2020), R core functions and the ‘poppr’ package (Kamvar *et al*. 2015).

The clustering tendency of the genotype matrix was estimated using the approach of Lawson and Jurs (1990). We clustered the LD-pruned genotype matrix using the following clustering algorithms: k-means, partition around medoids (PAM), clustering large applications (CLARA) and hierarchical clustering. In each algorithm, we allowed 2 through 30 clusters. Cluster quality was assessed using connectivity (Handl *et al*. 2005), Dunn index (Dunn 1974), and silhouette width (Rousseeuw 1987).

### Experimental layout and phenotyping

The TDP-300 was grown in the field in 2 consecutive years under well-watered (WW) and water-limited (WL) conditions. In 2020, the experiment was conducted at the J. Margulies Experimental Farm of the Hebrew University of Jerusalem in Rehovot (31.906° N, 34.799° E), Israel, using an augmented experimental design with 17 check lines replicated in three blocks. In 2021, the experiment was conducted at the Kvutzat Shiller farm (31.879° N, 34.777° E), Israel, using a factorial (lines x treatment) split-plot design (irrigation in main plots, lines in subplots) with three replicates. Soil types at the experimental locations were sandy loam and clay, respectively. Both experiments were sown on 22 April of the respective years. Experimental plots in both years consisted of six 2-m long and 15-cm spaced rows, sown at a seed rate equivalent to 6 kg ha^-1^ based on our previous study and the common practice in Israel (Ben-Zeev et al. 2020b). Experiments were sprinkler-irrigated until 2 weeks after emergence (WAE), then drip-irrigated. Due to technical constraints, a total of 216 and 252 TDP-300 lines were tested in the 2020 and 2021 experiment, respectively; of these, a complete data set across all four environments (2 years x 2 treatments) was available for 202 lines. Mineral nutrition and weed control were applied according to commercial farming practices, and no other pesticides were required.

Lodging was assessed visually by two independent surveyors, once a week from 8 to 10 WAE, using a scoring method adapted from Caldicott and Nuttall (1979) and recently described in Ben-Zeev et al. (2020b). Lodging was scored on a 10-level severity scale (0 being an upright, non-lodging plant and 9 being a fully lodging plant), and lodging prevalence was assessed (percentage of the entire plot area). For each plot, the severity was multiplied by the prevalence to calculate the lodging index (LI).

### Data analyses and heritability estimates

All data were analyzed using JMP software (Version 15, SAS Institute Inc., Cary, NC, 1989–2021) and R 4.1.3 (R Core Team 2020). Line performance under each of the four environments: 2020 water-limited (20WL), 2020 well-watered (20WW), 2021 water-limited (21WL), and 2021 well-watered (21WW), was calculated according to the experimental designs employed in each year. Owing to the augmented experimental design in 2020, data of individual plots were adjusted for block effects within each treatment, based on the 17 check lines replicated in each block. For the 2021 experiments, least square (LS) means were calculated using a full factorial model including E x G interaction and blocks. In addition, LS means were calculated jointly for all four environments using a model consisting of environment and genotype factors.

Broad sense heritability (*h^2^*) estimates of LI in each week were calculated using two approaches. For the replicated 2021 experiment, *h^2^* was calculated using variance components estimated based on ANOVA:

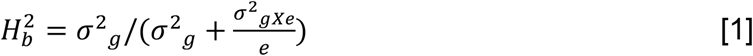

where *σ*^2^_g_ = [(MS _(G)_ - MS _(G × E)_/e], *σ^2^_gXe_* = [(MS _(G)_ + MS _(G × E)_/e], e is the number of environments and MS is the mean square.

Because ANOVA was not applicable for the augmented (non-replicated) 2020 experiment, and based on tef’s extreme self-pollination nature (Ketema 1983a), *h^2^* estimates of LI in each week were calculated using a linear regression coefficient (Cahaner and Hillel 1980) between the two treatments within the same year. Similarly, *h^2^* was calculated across years (2020 vs. 2021) for each treatment as well as for means of each year.

### Genome-wide association

Genotypic and phenotypic data were combined to perform a genome-wide association analysis using a mixed linear model including the full VCF file and an IBS kinship matrix. Due to the low clustering tendency of the population (see Results section), principal components were not included in the final model.

The threshold for significance was set based on the Bonferroni correction (0.05/n = 1.7 x 10^-6^, where n is the number of markers) and the Benjamini-Hochberg method implemented in the R package ‘RAINBOWR’ (Bonferroni 1936; Benjamini and Hochberg 1995; Hamazaki and Iwata 2020; R Core Team 2020). Manhattan plots and Q-Q plots were also produced using R. The association test was conducted using TASSEL (Bradbury *et al*. 2007). The mixed linear model, which included an IBS kinship matrix was:

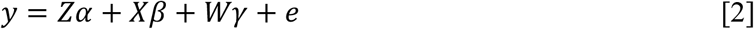

where y is the phenotype value, *α* is the value of the fixed effect other than SNP, *β* is the vector of the SNP effect, *γ* is the vector of the kinship effect, and e is the vector of residual effects.

## Results

### Sequencing quality

The final genotypic data included 28,837 SNPs, representing an average density of 46 SNPs/Mbp. The LD-pruned data set consisted of 9,268 SNPs. Nevertheless, use of the unpruned data was preferred to enable the detection of all significant SNPs. Density of SNPs varied along the genome, but no large regions devoid of SNPs were observed (Fig. S2). Missing data per site and per sample were at a low level, with ~10% at the third quartile (Fig. S1a). Most sites (27,550) were completely homozygous, and only 146 sites displayed >10% heterozygous genotypes (Fig. S1b). A quarter of all loci displayed MAF < 0.05 indicating an excess of rare alleles (Fig. S1c). The nucleotide diversity (π) exhibited a dichotomous distribution, skewed toward its possible extreme values of 0 and 0.5; the left skew is in line with the high abundance of rare alleles, whereas the right skew could be the result of (almost) fixed alleles between the two subpopulations (Fig. S1d).

### No subpopulation structure found

Estimated clustering tendency was 0.66, indicating nearly random data distribution. The cross-entropy criterion of sNMF was equivocal (Fig. S3). Possible candidates for the optimum number of K were 8, 14, and 20. The optimum number of clusters using classical clustering algorithms (k-means, PAM, CLARA, hierarchical) depended on the method employed for cluster-quality assessment. Connectivity, Dunn index, and silhouette width suggested 2, 2, and 30 clusters, respectively. Based on the two former clustering methods (out of the three), two subpopulations seemed to best describe the data. Due to the rather piecemeal data available regarding collection sites and environmental backgrounds for the TDP-300 lines, we could not attribute the clustering to any underlying parameter. Yet, in line with the low clustering tendency value, a clear-cut subpopulation structure was not observed, and the optimum number of clusters could not be unambiguously determined. Consequently, almost all samples exhibited at least some degree of admixture regardless of the number of clusters, and admixture increased with the growing number of subpopulations (Fig. S3). PCoA results corroborated the structure found using clustering algorithms (Fig. 2).

**Figure 2.**
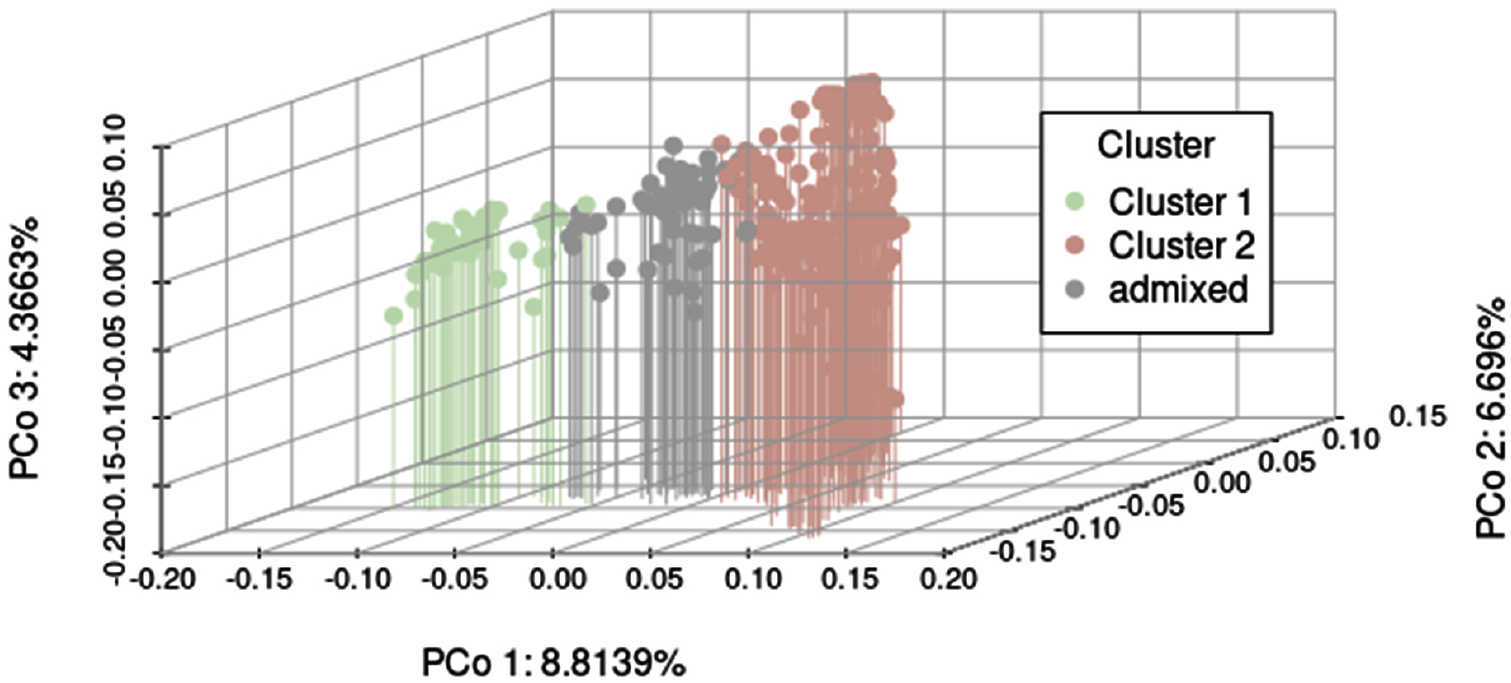
Genomic Principal coordinate analysis (PcoA) showing the relationship between the two subpopulations. Principal component (PC) 1 and 2 cumulatively explained ~15.5% of the overall genetic variation

### Dynamics of lodging in tef

Wide ranges of LI values were observed within each environment across various sampling times (Fig. 3). In both years, lodging onset was recorded at 8 WAE, as reflected by the distribution of LI values at this time point: low LI values for most lines while a few lines exhibited severe lodging, and these shifted to higher values in subsequent weeks. The natural trend of lodging escalation was evident from the increase in average values in any of the four environments along the presented 3-week data. As a result, LI did not distribute normally in any environment or at any sampling time and various transformations failed to restore normality. Therefore, LI data recorded at 8 WAE, which were heavily skewed toward lower values, were not used for heritability or GWAS analyses.

**Figure 3.**
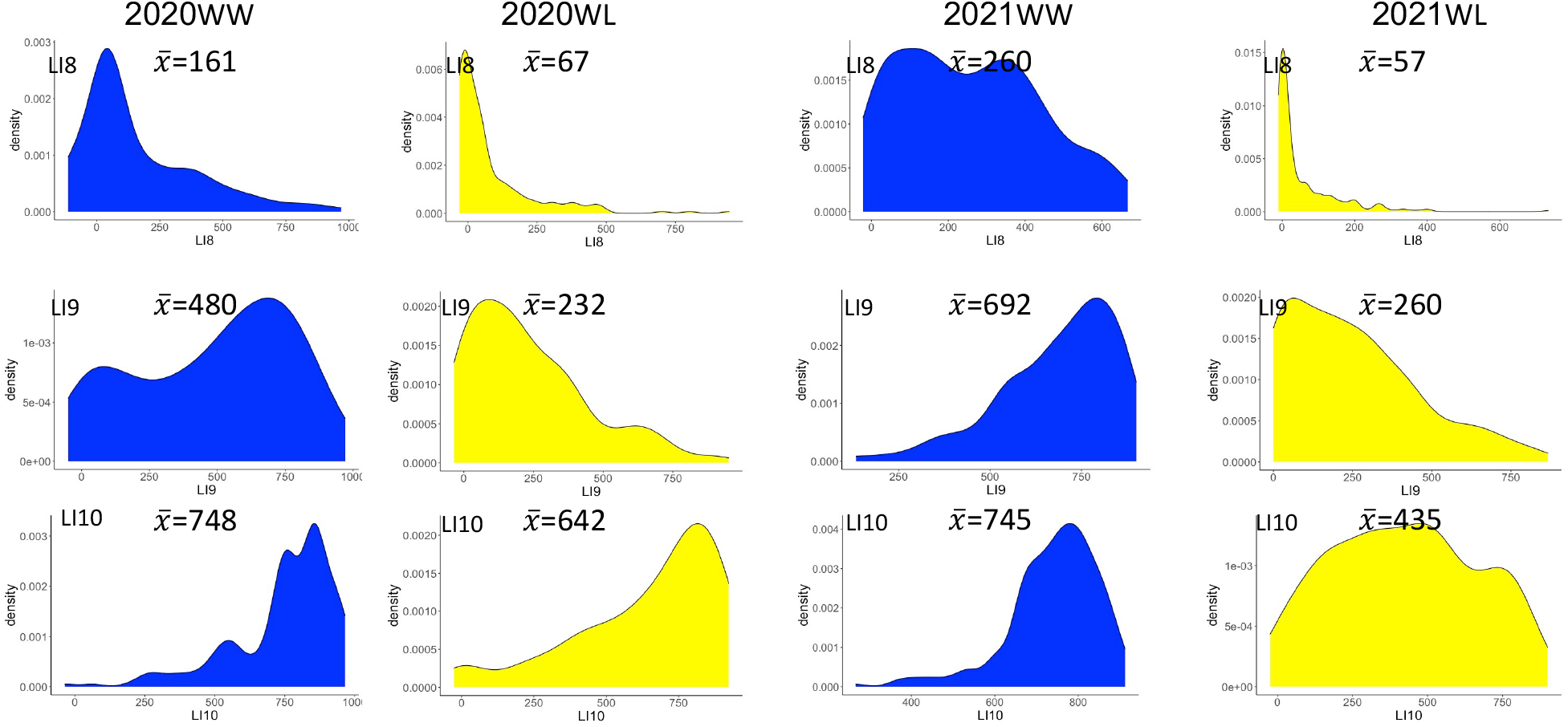
Distributions of lodging index values at 8, 9 and 10 weeks after emergence (LI8, 9, 10, respectively) under well-watered (WW) and water-limited (WL) treatments in the 2020 and 2021 seasons.

Despite being drip-irrigated, where water does not impose physical force on the crop’s canopy, greater LI values were recorded in the WW vs. WL treatment, with an up to 5-fold difference at 8 WAE in the 2021 season (Fig. 3). Notably, the WL treatment resulted in later lodging onset than the WW treatment in both years.

### Lodging has low to medium heritability

ANOVA-based *h^2^* estimates for LI (Eq. 1) using the 2021 data set were 0.27–0.30 (Table 1). Correlation-based *h^2^,* calculated for the 2020 experiments, resulted in higher values (0.49–0.55), while between-year estimates were intermediate (0.26–0.49) (all correlation coefficients were significant, *P* < 0.0001). Owing to the self-pollinating nature of tef, all loci are expected to be homozygous and with no dominant effect. Therefore, correlation-based *h^2^* is a good proxy for the additive variance.

**Table 1.** ANOVA-based and correlation-based broad sense heritability (h^2^) estimates of lodging indices at 9 and 10 weeks after emergence (LI9 and LI10, respectively) and water-limited (WL), well-watered (WW) or averages across environments (LS means).

| Data sets used | $h^2$ estimates | |
| --- | --- | --- |
|  | LI9 | LI10 |
| ANOVA-based estimate |  |  |
| 2021 (two environments) | 0.30 | 0.27 |
| Correlation-based estimate |  |  |
| 2020WL vs. 2020WW | 0.49 | 0.55 |
| 2020WL vs. 2021WL | 0.44 | 0.48 |
| 2020WW vs. 2021WW | 0.26 | 0.36 |
| LS means 2020 vs. LS means 2021 | 0.44 | 0.49 |

### Twenty nine sites found to be associated with lodging

A GWAS was conducted for eight LI data sets separately, each representing a specific year (2020, 2021), treatment (WW, WL) and time (9 or 10 WAE) (Figs. 4, 5). Manhattan plots for each data set are presented with Bonferroni-corrected threshold and Benjamini-Hochberg threshold (applicable only when the former threshold was met).

**Figure 4.**
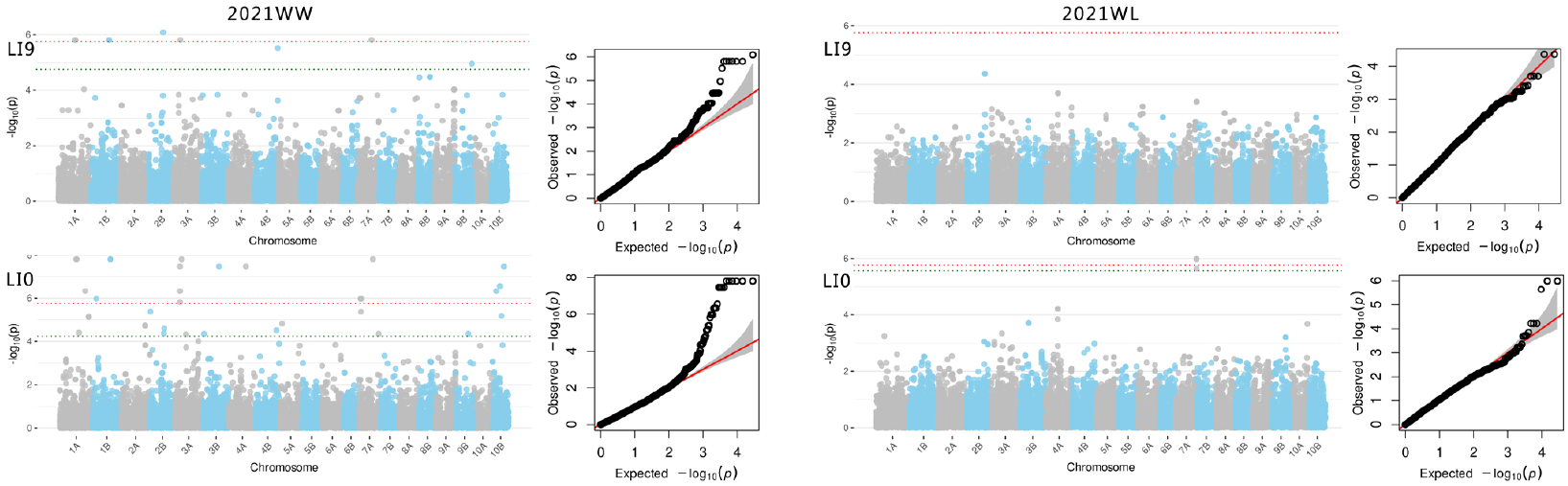
Manhattan and Q-Q plots of the association of 28,837 SNP with lodging index at 9 and 10 weeks after emergence (LI9 and LI10, respectively) under well-watered (WW) and water-limited (WL) treatments in the 2020 season. Red and green dotted lines (respectively) represent the significance threshold based on the Bonferroni correction threshold and Benjamini-Hochberg correction threshold (where applicable).

**Figure 5.**
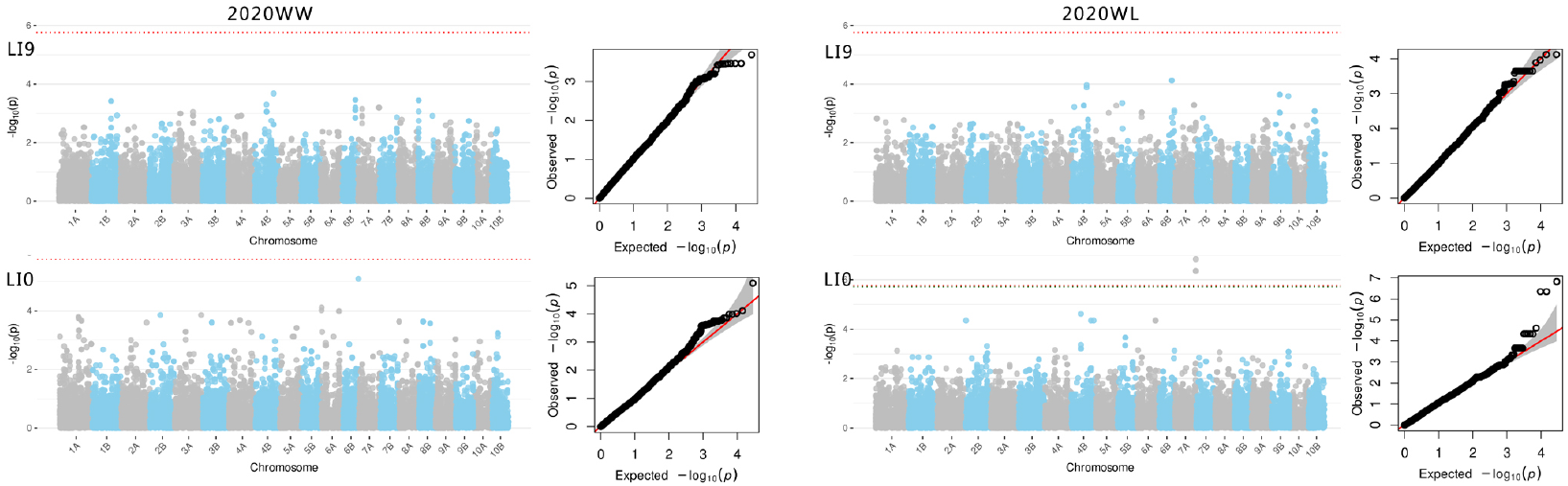
Manhattan and Q-Q plots of the association of 28,837 SNP with lodging index at 9 and 10 weeks after emergence (LI9 and LI10, respectively) under well-watered (WW) and water-limited (WL) treatments in the 2021 season. Red and green dotted lines (respectively) represent the significance threshold based on the Bonferroni correction threshold and Benjamini-Hochberg correction threshold (where applicable).

A two-step approach was employed to select the most important sites associated with LI. First, we screened for significant sites (meeting the Bonferroni-corrected threshold, 1.7 x 10^-6^) or a less stringent suggestive threshold (*P* < 1 x 10^-4^) determined based on lowest *P-*values indicated by the Benjamini-Hochberg correction. This process resulted in a list of 29 sites distributed across 10 of the 20 tef chromosomes (Table S1). The number of lines carrying the minor alleles varied between 3 and 22 across the 29 significant or suggestive sites.

To minimize the potential of false positives, we then selected sites that were significant (Bonferroni threshold) in at least one data set and suggestive in at least one other data set. This process resulted in 11 selected sites of prime interest. These 11 sites outlined four groups of lines which shared the same minor allele(s) at common site(s) (Table 2). Group 1 consisted of four TDP-300 lines with the same minor alleles at six common sites (on chromosomes 1A, 1B, 3A and 7A). Groups 2 and 3 consisted of seven and eight lines, respectively, having the same minor allele at a single site each (chromosome 2B and 1A, respectively). Importantly, there was some overlap between lines belonging to groups 1–3. Group 4 consisted of 21 lines with the same minor allele at three common sites (all three on chromosome 7A). The lines in group 4 were not present in groups 1 and 2.

**Table 2.**
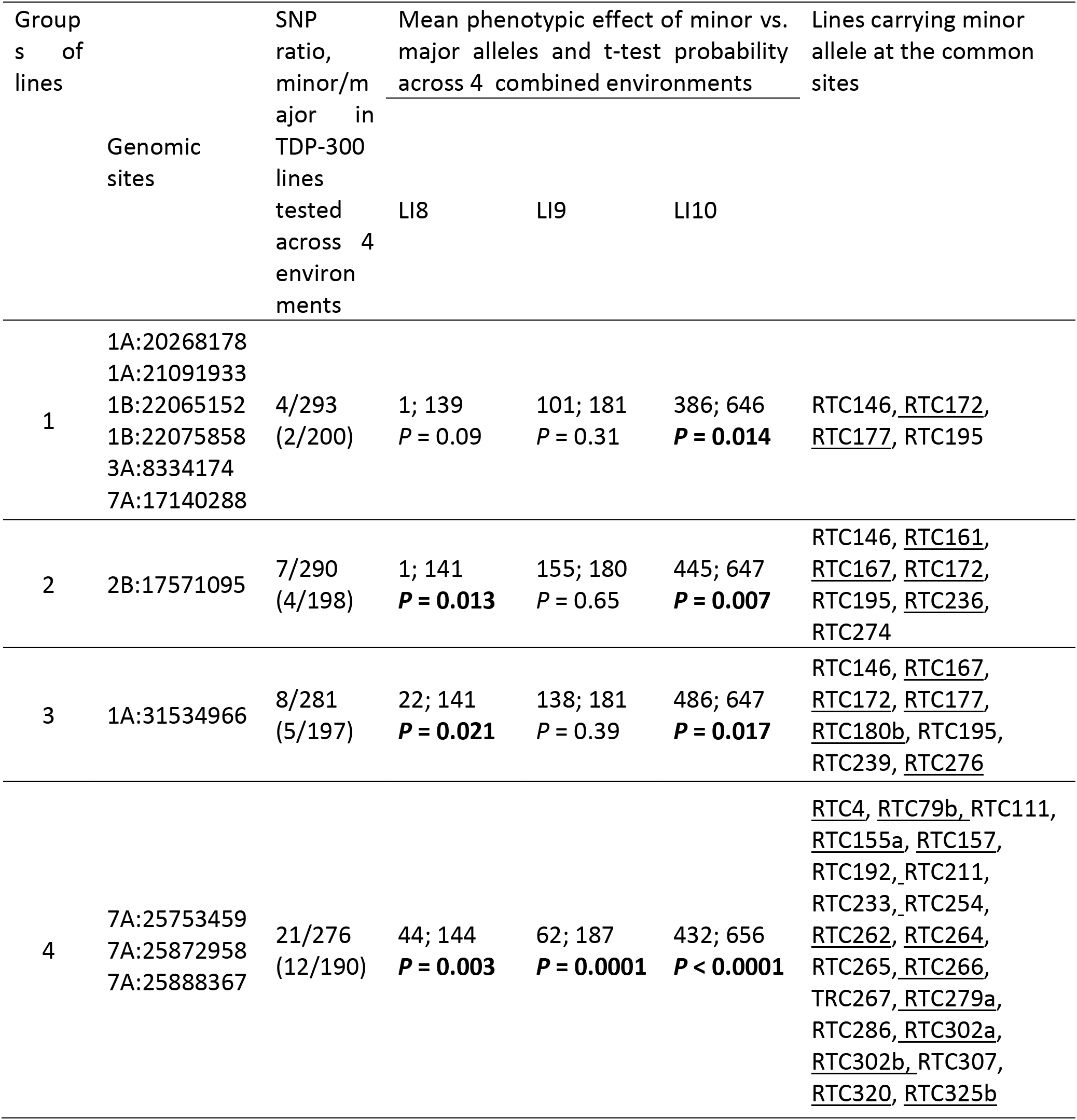
Selected genomic sites associated with lodging indices (LI) in two or more data sets (see Table S1) divided into four groups carrying a minor allele at the common sites of interest. Presented are SNP ratio in the TDP-300 lines tested across four environments (out of 202), mean LI of the various allelic groups and *P*-values of t-test for LS (significant *P-values* in bold) means for all four environments combined. Lines carrying minor allele are listed; those tested across all environments are underlined.

A t-test was used to test the effects of selected sites on LI recorded at 8-10 WAE, averaged (LS means) across all four environments (Table 2). The phenotypic differences between the two allelic groups were significant in most cases (*P* < 0.05), thus validating the effect of the selected sites. The difference between lines carrying the minor vs. major alleles ranged between 100 and 280 LI units, nearly one-third of the entire LI range.

The tef genome browser (genomevolution.org/coge/GenomeInfo.pl?gid=50954) was used to search for candidate genes located at or closest to (up to 200 kbp up or downstream) the selected genomic loci associated with LI. Out of the 11 selected sites, 4 were located in genes (all 4 with a putative function) and 7 were located near genes (1 with a putative function) (Table 3).

**Table 3.** Candidate genes located at (in bold) or near (200 kbp up or downstream) the selected genomic sites associated with LI.

| Genomic site | Genes at site |
| --- | --- |
| 1A:20268178 | Uncharacterized |
| 1A:21091933 | Uncharacterized |
| 1B:22065152 | ***Setaria italica* acyl transferase 1 \*** |
| 1B:22075858 | Uncharacterized |
| 3A:8334174 | Uncharacterized |
| 7A:17140288 | Uncharacterized |
| 2B:17571095 | ***Panicum virgatum* LRR receptor-like serine/threonine-protein kinase RGI3 \*\*** |
| 1A:31534966 | ***Panicum virgatum* cilia- and flagella-associated protein 58-like** |
| 7A:25753459 | ***Setaria viridis* glyoxylate/hydroxypyruvate reductase HPR3-like** |
| 7A:25872958 | Uncharacterized |
| 7A:25888367 | ***Setaria viridis* pentatricopeptide repeat-containing protein \*\*\*** |

\* Acyltransferase was associated with suberin synthesis in roots and seed coats (Beisson *et al.*, 2007)
\*\* A gene family expressed in various parts of the roots (Wu *et al.*, 2016)
\*\*\* PPR proteins were involved in root and shoot development and abiotic stress (Lee *et al.*, 2019; Su *et al.*, 2019)

## Discussion

### TDP 300 -a new resource for tef genomics

Being a minor crop with narrow geographic distribution, availability of genomic infrastructure, genetic markers, and a mapping database is limited for tef (Tadele 2019). Hence advanced genome-based, marker-assisted breeding of tef has remained mostly out of reach thus far (Bekana and Assefa 2021; Numan *et al*. 2021). Genomic sites related to lodging have scarcely been reported in tef, although a few studies pre-dating the published tef genomes (Cannarozzi *et al*. 2014; VanBuren *et al*. 2020) reported on genetic markers associated with lodging (Yu *et al*. 2007; Zeid *et al*. 2012). The difficulty involved in crossing tef (Berhe 1975; Zeid *et al*. 2010; Habteab *et al*. 2015) and the deficiency of reference genomes (Adnew *et al*. 2005; Tadele 2019; Girija *et al*. 2021) may have hindered the development of biparental populations and diversity panels, respectively, both being essential for genetic mapping. Hence, the TDP-300 reported in the current study may become a useful tool for genomic dissection of various traits of interest in tef and subsequently, their advancement via marker-assisted breeding.

To test the clustering tendency, Lawson and Jurs (1990) compared the observed structure with the structure of “white noise”. In the absence of any clusters, the expected value was 0.5, whereas a value of 0.75 indicated a clustering tendency at the 90% confidence level (Han *et al*. 2011). In the TDP-300, the estimated tendency was 0.66, suggesting very low clustering. In line with this, the clustering approaches produced ambiguous results and did not indicate a single optimum number of clusters that would be favorable for GWAS. We opted to use two clusters (subpopulations) as the optimum number of subpopulations to apply the coloring schemes in the figures (Figs. S3, S4), because two independent approaches pointed to this cluster number. Yet, the varying results indicated that the collection cannot be described by any clear-cut subpopulation structure. Hence, for GWAS, we assumed no subpopulation structure in the TDP-300.

### Phenotypic distribution suggest rare lodging resistance and strong environmental effect

Lodging is a complicated phenomenon resulting from plant (shoot and root) traits (Berry *et al*. 2004), environmental factors (soil properties and moisture) (Pinthus 1974; Berry *et al*. 2000), and management practices (irrigation, sowing depth and density) (Ben-Zeev, Kerzner, *et al*. 2020; Ben-Zeev, Rabinovitz, *et al*. 2020). The nature of tef lodging as a dynamic and largely irremediable phenomenon is clearly depicted in the LI distributions at 8–10 WAE across the four studied environments (Fig. 3). Due to the rather uniform onset of lodging and the rarity of lodging-resistant lines, LI distribution was heavily skewed toward lower values at 8 WAE, became wider at 9 WAE (Fig 1a), and moved toward the high end of LI values at 10 WAE.

Heritability estimates ranged between 0.26 and 0.55 (Table 1) at different times (WAE), under different environments, and with different calculation methods. In line with our results, previous studies on tef lodging have reported *h^2^* values of 0.16 to 0.55 (Chanyalew 2010; Ayalneh *et al*. 2011; Bekana and Assefa 2021). In the current study, lower values (0.27–0.30) were obtained from the ANOVA-based calculation, and the highest values from a between-environment correlation in the 2020 season. Since correlation-based *h^2^* tends to be overestimated, it is the authors’ opinion that *h^2^* values of 0.30–0.45 for tef lodging are probably realistic. These estimates, considered low to intermediate, reflect the small genetic variation and large effects of environment and management on tef lodging.

### Lodging related genomic sites group together and associate with root development genes

Owing to the nature of lodging in tef and the rarity (or near absence) of non-lodging lines, LI did not distribute normally, particularly at 8 WAE. Transformations failed to restore normality, and therefore LI8 was excluded and the original data sets of LI9 and LI10 were used for GWAS, albeit with a careful consideration of the results. A total of 29 significant (Bonferroni-corrected threshold) or suggestive (less stringent threshold) sites were detected in the current study (Table S1); in all cases the minor allele conferred reduced lodging, in line with the rarity of lodging resistance in tef. An additional screening resulted in 11 selected sites which outlined four groups of lines sharing the same minor allele(s) at common site(s) (Table 2). Groups 1 and 4 consisted of 6 and 3 common sites, respectively (Table 2). The current data and panel do not allow distinguishing between the effect of the various sites within each group, and their apparent phenotypic effect may result from either a single site (with all other sites being neutral), closely linked sites or a few independent sites. Previous mapping studies of tef lodging (Yu *et al*. 2007; Zeid *et al*. 2011) pre-dated the published reference genomes, and therefore we could not compare the previously published sites with those identified in the current study.

Due to the lodging-assessment method, LI ranged between 0 and 900. A somewhat wider distribution resulted from the block adjustment of the 2020 augmented design experiment or the LS mean calculated for the 2021 replicated experiment (Fig. 3). The effects of SNP sites on LS mean LIs of the four combined environments confirmed the effects of the selected sites and demonstrated the magnitude of the genetic effect on LI (Table 2). Various sites seem to affect lodging at different times during plant development, suggesting their involvement in different plant traits and/or developmental processes, such as crown root number and diameter, culm diameter and number of tillers, as suggested in our previous publication (Ben-Zeev et al. 2020b). A number of published studies have targeted reduced plant heightt as a means of reducing lodging in tef (Zhu et al. 2012; Jost et al. 2015; Jifar et al. 2017; Bekana and Assefa 2021). Mapping plant height and panicle length assessed under the same four environments identified a number of sites associated with these traits (data not shown). However, no overlap was found between sites associated with lodging and those associated with plant height or panicle length. While plant height determines the bending moment applied by the canopy, root-borne lodging depends also on the diameter of the root-soil cone, the bending strength of the crown and the shear strength of the soil (Shah *et al*. 2019).

Three of the five candidate genes with a putative function were reported to be involved in root development (Table 3). Genes found at or close to genomic sites associated with lodging belong to gene families which control suberin production in roots (Beisson *et al*. 2007), root development (Wu *et al*. 2016), and stress response (Lee *et al*. 2019; Su *et al*. 2019). The putative functions of genes identified demonstrate the important role of root traits in tef lodging, and in turn suggest new targets for lodging resistance breeding in tef.

### Concluding remarks

This study presents the first tef diversity panel (TDP-300) subjected to modern genotyping technique (GBS-based SNP markers). Phenotyping of TDP-300 under four environments in the field revealed wide diversity in a number of phenotypic traits, most of which will be presented in our follow-up publication. The current study demonstrates the utility of TDP-300 for genomic mapping of complex traits in tef. Seeds of TDP-300, as well as our VCF, are available for distribution to allow further research of this important and intriguing orphan crop, which in turn will support broader tef cultivation and production (Tadele 2019; Kebede 2021; Numan *et al*. 2021).

The sites identified in this study were found to reduce the LI by up to 280 units (Table 2), which may play a vital role in reducing lodging damage in traditional as well as modern tef cultivation, thus improving farmers’ livelihood and food security. Combining several alleles associated with lodging resistance may result in a greater reduction of LI. The identification of lodging-related sites in the current study, as well as in future studies, can help understand the mechanisms underlying tef lodging and provide a basis for advanced genome-based, marker-assisted breeding of tef

## Abbreviations

GWAS: Genome wide association study
LI: lodging index
SNP: Single nucleotide polymorphism
TDP-300: tef diversity panel 300
VCF: variant calling file
WAE: weeks after emergence
WL: water-limited
WW: well-watered

## Acknowledgements

This study was supported by The Israel Ministry of Agriculture and Rural Development, Chief Scientist Foundation [grant no. 12-02-0008] and The Israel Innovation Authority, Challenge Program [grant no. 73546]. The authors wish to acknowledge the use of the services and facilities of AGRF. We thank the Netafim irrigation company for providing the irrigation system for the 2020 season. We are grateful to Noa Kirby, Philip Wagali, Ido Zeltzer, Neta Levinson, Ori Harash, Ariel Joseph, May Shoshan, Eran Dagan, Emma Braslavsky, Eric and Rotem Ben-Zeev who enabled performance of this study with their hard work and insights. S.B.-Z. is indebted to the Robert H. Smith Foundation for a doctoral fellowship award. M. D. is The Heinrich Bonnenberg Scholarship Awardee. Y.S. is the incumbent of the Haim Gvati Chair in Agriculture.

## Author Contributions

Shiran Ben-Zeev and Yehoshua Saranga assembled the diversity panel, conceptualized and designed the study. Shiran Ben-Zeev Muluken Demeile and Vered Barak collected the data, propagated the plants and extracted the DNA. Timo Hellwig conducted the population structure and genomic data analysis and supported the association study. Sasha Vorobyova and Sariel Hübner analyzed the raw genomic data and provided insight into the population structure and the association study. Shiran Ben-Zeev and Yehoshua Saranga analyzed the phenotypic and genomic data and wrote the first draft of the manuscript. All authors read and approved the final manuscript.

## Data availability statement

Genomic data and seeds are readily available for the scientific community. VCF files will be made available on our web site, sequence data as well as seeds of the TDP300 will be provided upon request.

## Supplementary Figures

**Figure S1.**
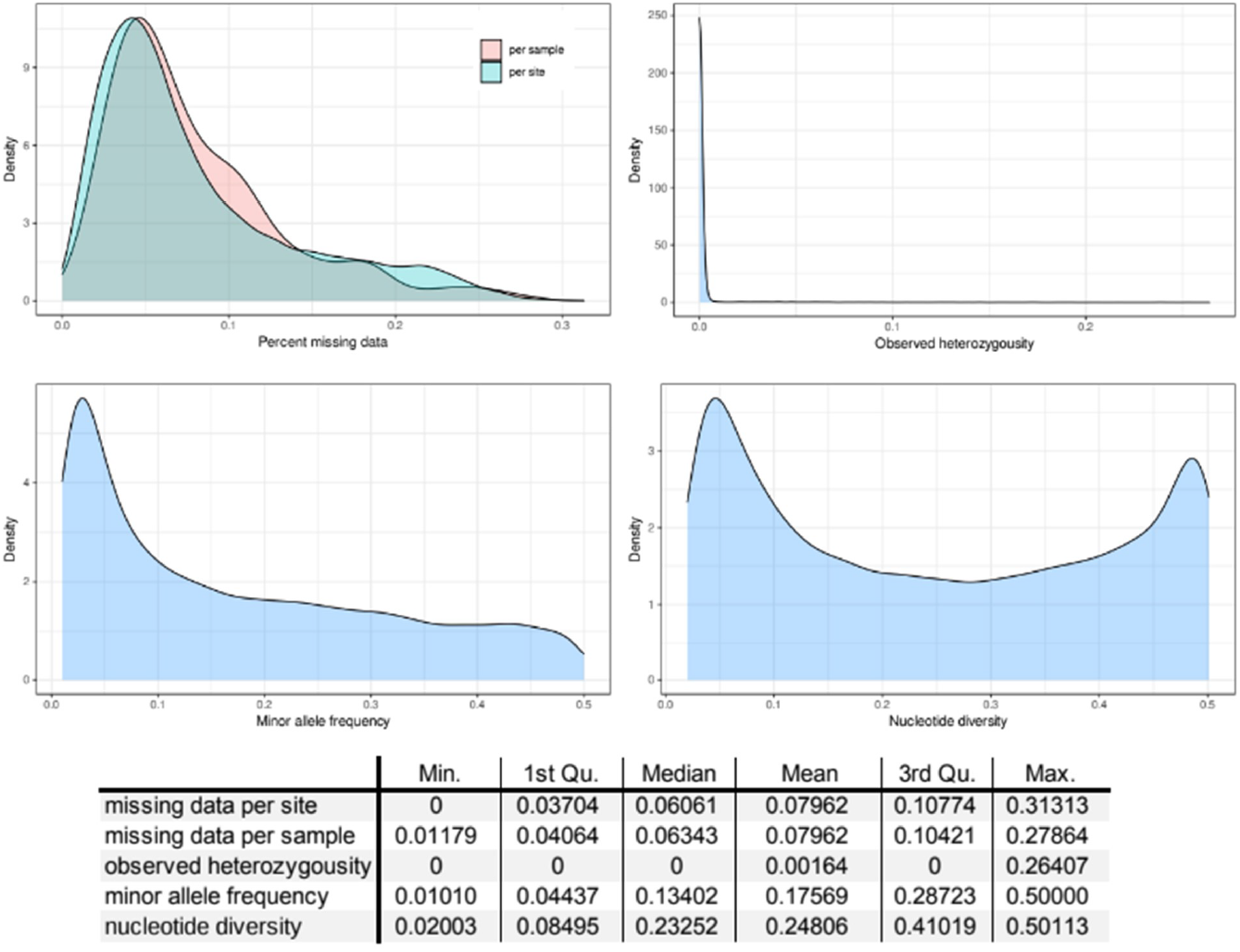
Quality assessment of the final SNP set described by missing data per site and sample, observed heterozygosity, minor allele frequency and nucleotide diversity (π). Min., minimum; 1st Qu., first quartile; 3rd Qu., third quartile; Max., maximum.

**Figure S2.**
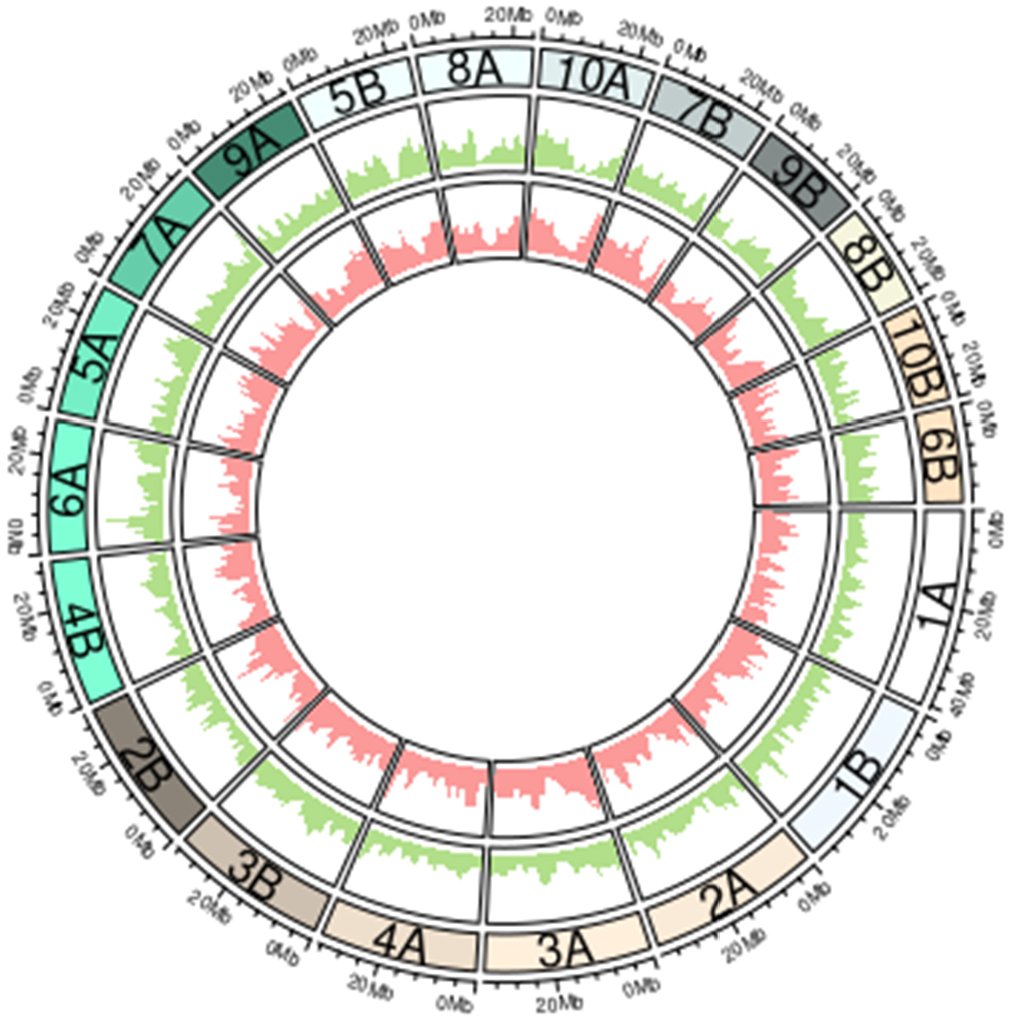
Distribution of SNP density over chromosomes. Outer circle (green): filtered variant set. Inner circle (red): LD-pruned variant set.

**Figure S3.**
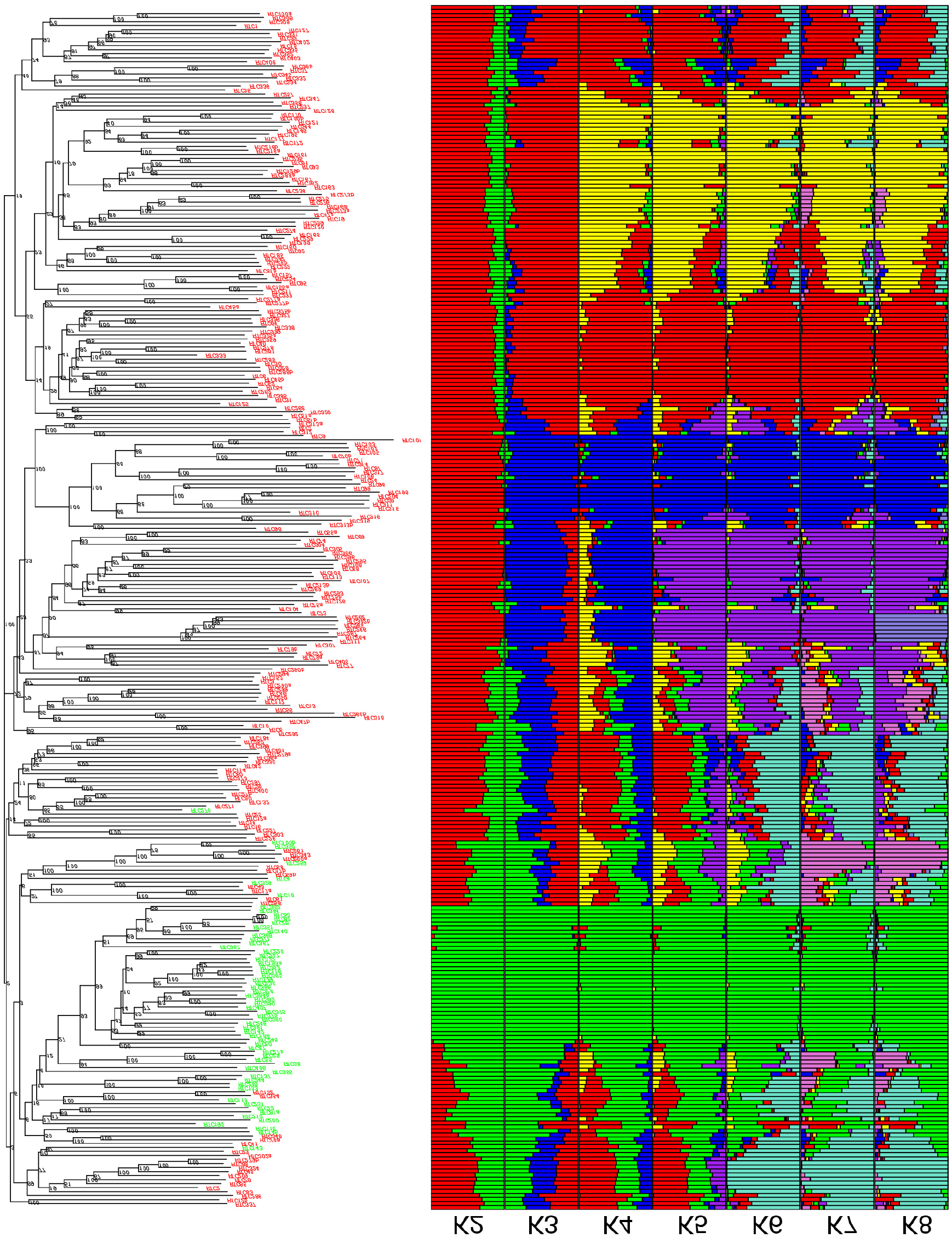
Neighbor-joining tree and sNMF bar plots with the number of subpopulations, K, ranging from 2 to 8. Only bootstrap values >70% are shown. Sample labels in the neighbor-joining tree are colored according to sNMF subpopulation assignment, with K = 2 if they had at least 60% subpopulation fraction of one subpopulation.

**Figure S4.**
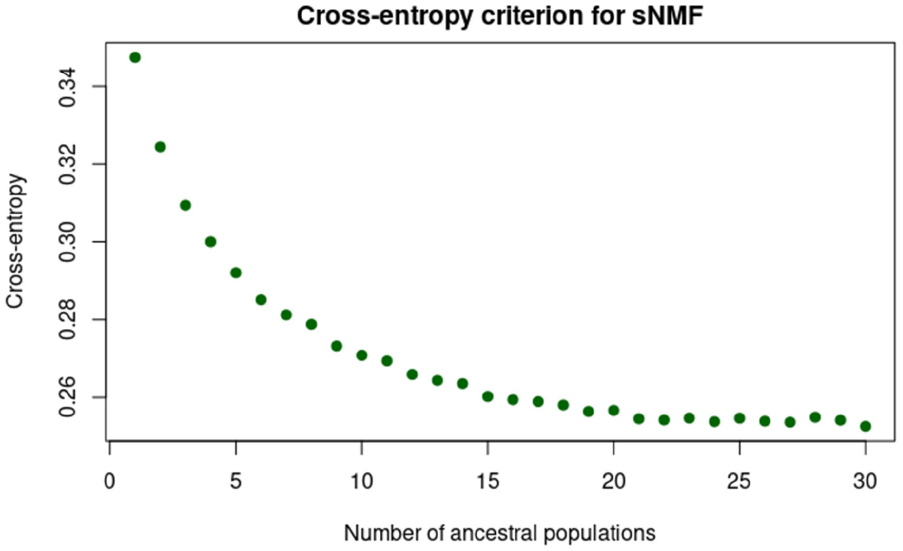
Cross-entropy criterion of sNMF averaged over replications for K ranging from 0 to 30.

---

**Table S1.**
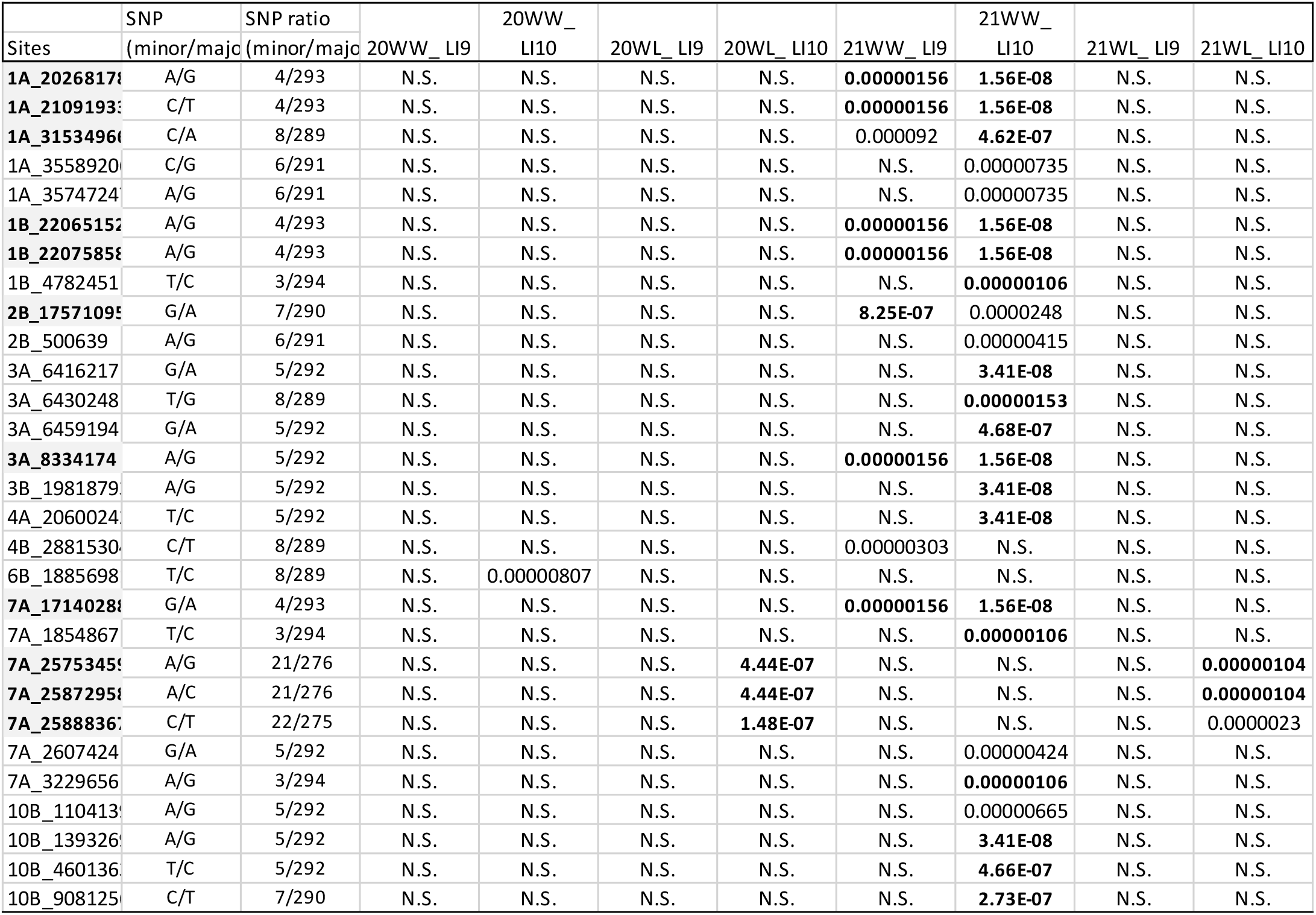
significant sites (Meeting the Bonferroni threshold, 1.7 × 10-6), and suggestive threshold (p<1*10-4)

